# The HSFA1a is stabilized by the co-chaperone HSP70-HSP90 organizing protein HOP in Arabidopsis

**DOI:** 10.1101/2024.01.30.577911

**Authors:** René Toribio, Aurora Navarro, M. Mar Castellano

## Abstract

Living organisms have the capacity to sense and respond to environmental stimuli, including warm conditions. Upon the increase of temperature, eukaryotic cells promote the onset of a universal transcriptional response called the heat stress response (HSR). HSR is triggered by the activation of the heat shock factor 1 (HSF1), whose activity is highly regulated in all eukaryotes. A common process of HSFA1 regulation involves its binding to HSP70 and HSP90 chaperones. However, the effect of this binding differs among eukaryotes and is not fully elucidated in plants. Moreover, the role of chaperone auxiliary proteins, such as HOP, in HSF1 regulation remains unknown. Here, we show that HOP is involved in the folding and stabilization of the HSFA1a and is required for the onset of the HSR during thermomorphogenesis in Arabidopsis. Our results demonstrate that the members of the AtHOP family bind in vivo to the HSFA1a and that the expression of multiple genes involved in the HSR, including numerous HSFA1a-responsive genes, is altered in the *hop1 hop2 hop3* mutant under warm temperature. HOP is not involved in the subcellular localization of the AtHSFA1a. Instead, HSFA1a is accumulated at lower levels in the *hop1 hop2 hop3* mutant while control levels are recovered in the presence of the proteasome inhibitor MG132 or the synthetic chaperone TUDCA. This uncovers the HSFA1a as a new client of HOP in plants and reveals the participation of HOP in HSFA1a stability, expanding the notorious function of the co-chaperone HOP in thermomorphogenesis.

## INTRODUCTION

One effect associated with climate change is the increase of the global temperature of the Earth (also known as global warming). Current climate models estimate that the global average temperature will rise 1-5°C at the end of the 21^st^ century (Driedonks et al., 2016). This moderate increase in temperature will lead to drastic losses in the production of important crops as wheat, rice or corn in tropical and subtropical areas (Challinor et al., 2014; FAO, 2009), which would highly limit the main source of food worldwide.

As sessile organisms, plants are forced to cope with and adapt to the environmental challenges, including mild heat. In this sense, it is well known that plants have evolved different mechanisms to allow their growth when ambient temperature exceeds a few Celsius degrees over their optimum level. One of these mechanisms involves changes in plant morphology. These changes aim to preserve the meristems and favour transpiration to avoid the harmful effects of heat. Such morphology changes, globally known as thermomorphogenesis, include an increased elongation of the hypocotyl, roots and petioles from rosette leaves and cotyledons (Quint et al., 2016).

Thermomorphogenesis is a complex process highly regulated at multiple molecular levels, including a sophisticated regulation of gene expression. In such a way, thermomorphogenesiś transcriptional output results from the coordinated action of multiple signaling pathways. These include the activation of different hormonal networks, such as auxins, gibberellins (GAs) and brassinosteroids (BRs). In addition, it relies on the activation of PHYTOCHROME INTERACTING FACTOR4 (PIF4), a transcription factor that has a large level of crosstalk with the hormonal networks (Cortijo et al., 2017; Quint *et al*., 2016). Moreover, during thermomorphogenesis, plants also launch the so called heat stress response (HSR), which involves the up-regulation, among others, of the heat shock proteins (HSPs) (Cortijo *et al*., 2017). This transcriptional response is controlled by different heat-shock factors (HSFs), among which the HSFA1s are considered the master regulators, because they are the first to be activated and induce the downstream transcriptional response (Yoshida et al., 2011). Interestingly, the number of members of the HSFA1 family greatly differs among plants. While there is a single member of the HSFA1 family in tomato, four HSFA1s have been described in Arabidopsis (HSFA1a, b, d, e) (Yoshida *et al*., 2011). Among them, the AtHSFA1a has been specifically involved in the thermomorphogenesis response, since it binds, under conditions of mild increases of temperature, to the promoters of many HSR-responsive genes (Cortijo *et al*., 2017).

Since thermomorphogenesis relies on the activation of different hormonal and transcriptional networks, it is easy to visualize that it highly depends on the accurate accumulation and functionality of those proteins (such as receptors, transcription factors, kinases, etc.) that assure the perception and the correct signaling of the response. In such a way, processes, such as protein folding, that directly impact on protein accumulation, stability or functionality are expected to play a relevant role in this response (Moran Luengo et al., 2018; Moran Luengo et al., 2019).

One important chaperone that promotes the folding of crucial signaling proteins in mammals and yeast is HSP90. In these eukaryotes, HSP90 remodels more than 200 proteins involved in a wide range of physiological processes, including cell cycle progression, signal transduction and transcriptional regulation (Kravats et al., 2018; Verma et al., 2016). In plants, the number of the proteins known to be folded by HSP90 (known as clients) is extremely low compared to other eukaryotes. Nevertheless, HSP90 has already been described as important regulatory node in JA and auxin hormonal signaling (Wang et al., 2016). Remarkably, associated to its role in auxin signaling, HSP90 has been involved in thermomorphogenesis. In this regard, it was demonstrated that the high accumulation of HSP90 in response to mild temperature enhances the folding and stability of the auxin co-receptors TIR1 and AFB2. In this way, by promoting the accumulation of these co-receptors, HSP90 participates in the efficient auxin signaling needed for the correct establishment of the thermomorphogenesis response (Wang *et al*., 2016).

HSP90 is usually assisted in the folding by the chaperone HSP70 and by a cohort of co-chaperones, which regulate HSP90 specificity and function {di Donato, 2019 #893. One of these co-chaperones is the HSP70-HSP90 organizing protein (HOP). HOP binds simultaneously HSP70 and HSP90 and facilitates the transfer of the client proteins from the chaperone HSP70 to the HSP90 (Baindur-Hudson et al., 2015; Bhattacharya and Picard, 2021; Johnson et al., 1998; Lee et al., 2012; Odunuga et al., 2004; Toribio et al., 2020). In contrast to mammals and yeast that have only one *HOP* gene, the number of *HOP* genes codified by the plant genomes differs among species. In this way, Arabidopsis HOP family encompasses three different members HOP1, HOP2 and HOP3 Interestingly, these proteins have been proposed as master assistant of thermomorphogenesis (Castellano, 2024). Indeed, it has been recently described that AtHOP proteins contribute to hypocotyl elongation by assuring the proper accumulation of the F-box proteins TIR1 and SNE, and, therefore, an efficient auxin and GA signaling under warm temperatures (Mangano et al., 2023; Munoz et al., 2022). Although we already know some aspects of the role of HOP in this process, we still do not know whether the function of HOP in thermomorphogenesis is strictly circumscribed to the stabilization of these two important F-box proteins or if, apart from TIR1 and SNE, there could be other clients of HOPs involved in this adaptative response.

Here, we describe that HOP participate in the proper establishment of the HSR during thermomorphogenesis through the binding and folding of the HSFA1a, allowing the proper accumulation of HSFA1a under control and warm temperatures.

## MATERIALS AND METHODS

### Plant material and growth conditions

Arabidopsis T-DNA insertion mutants *hopl-l* (GK-420A10.15), *hop2-l* (GK-399G03.03), *hop3-1* (Salk_00794) and *hsfa1a* (Salk_125881) were acquired from the Arabidopsis Biological Resource Center (ABRC). The *hop1 hop2 hop3* triple mutant was previously described (Fernandez-Bautista et al., 2018).

Except otherwise stated, seeds were surface-sterilized, stratified at 4°C for 48 h and grown at 22°C using a 16-h light photoperiod. For in vitro growth, Murashige-Skoog (MS) medium supplemented with 1% (w/v) sucrose was used in all cases.

### Constructs and molecular cloning

To obtain the *pHSFAla:HSFAla-GFP* construct, an *AtHSFA1a* (AT4G17750) genomic fragment (-1597-1634 pb) was cloned into the binary vector *pGWB4* (Nakagawa et al., 2007). This construct was used to transform Col-0 and the *hop1 hop2 hop3* plants by the floral dip method (Clough and Bent, 1998). Homozygous lines were screened by RT-qPCR and 2 pair of lines that show similar levels of the *HSFA1a-GFP* transgene in Col-0 and *hop1 hop2 hop3* mutant within each line, were selected for further analyses.

The constructs *p35S:Flag-HOP1*, *p35S:Flag-HOP2* and *p35S:Flag-HOP3* were obtained by cloning the corresponding cDNA sequences in frame with the epitopes Flag in the *pGWBl2* binary vector (Nakagawa et al., 2007). In the case of the construct *p35S:HSFA1a-HA*, the genomic fragment of *HSFA1a* described above was cloned in frame with the HA epitope in the *pGWB4l4* binary vector. To generate the constructs for BiFC assays, the cDNAs of *HOP1*, *HOP2* and *HOP3* were cloned in frame with the N-terminal fragment of the Yellow Fluorescent Protein (YFP) while the genomic fragment of *HSFA1a* was cloned in frame with the C-terminal fragment of Cyan Fluorescent Protein (CFP) into the binary vectors pNXGW and pCXGW, respectively (Kim et al., 2009). pCXGW-tGUS was kindly provided by Dr. Guerrero’s lab, from the CBGP, and the *p35S:HOP3-GFP* was previously described in (Fernandez-Bautista *et al*., 2018). To obtain the *p35S:HSFA1a–mCherry* construct, the genomic fragment of *HSFA1a* was cloned in frame with the protein mCherry into a modified ppZP binary vector.

### Immunoprecipitation analysis and western blot

*N. benthamiana* leaves were agroinfiltrated with the corresponding constructs and harvested 3 days after the agroinfiltration. Immunoprecipitation analyses were carried out as described in (Toribio et al., 2019). Experiments were done at least three times, obtaining similar results. A representative replicate is shown in the figures.

For western blots, proteins were separated by SDS-polyacrylamide gel electrophoresis, blotted to nitrocellulose membranes and analyzed with specific antibodies. The monoclonal anti-Flag (Sigma-Aldrich), anti-GFP (Roche), anti-HA (high affinity clone 3F-10, Roche) and monoclonal anti-Actin (clone 10B 3, Sigma-Aldrich) were used with their respective HRP-secondary antibodies. Enhanced chemiluminescent reagent (GE Healthcare) was used to detect the proteins. Experiments were done at least three times, obtaining similar results. A representative replicate is shown in the figures. Quantification of protein accumulation in western blots was performed using the software ImageJ (https://imagej.nih.gov/ij/index.html). Statistically significant differences were calculated using the tests described in the figure legends.

### Microscopy analyses in Arabidopsis roots and quantification of HSFA1a-GFP accumulation

Roots from 7-days-old Arabidopsis seedlings of the *pHSFA1a:HSFA1a-GFP* lines (Col-0 and *hop1 hop2 hop3* mutant) that constitute the pair #1 and pair #2 were imaged at different temperatures and conditions as described in the figure legends. Experiments were done at least three times, obtaining similar results. A representative replicate is shown in the figures. After image capture, the intensity of nuclear HSFA1a-GFP in individual cells was analyzed and quantified using the image processing toolkit ImageJ. Data were normalized by computing the average values obtained from control conditions. Statistically significant differences were calculated using the tests described in the figure legends.

### BiFC and co-localization analyses

Leaves from 3-weeks-old *N. benthamiana* were transformed by agroinfiltration with different constructs to transiently express specific combinations of proteins (as stated in the figures or figure legends). Leaf discs were imaged 3 days post-agroinfiltration. In the case of BiFC, the constructs pNXGW and pCXGW-tGUS, which express the Nt-YFP moiety alone and the Ct-CFP moiety fused to a fragment of the unrelated protein *β*-galactosidase (GUS), were used along the constructs under study as negative controls. Images were captured using a Zeiss LSM 880 confocal microscope (Leica Microsystems) with an Argon ion laser. GFP was excited at 488 nm, and the emitted light was captured at 494–566 nm. Cherry was excited using 561 nm and emitted light captured at 578-618 nm. Sequential scanning was used to image GFP and Cherry as described in (Fernandez-Bautista *et al*., 2018). Experiments were done at least three times, obtaining similar results. A representative replicate is shown in the figures.

### RT-qPCR analyses

RT-qPCRs were performed as described in Echevarria-Zomeno *et al*. (2015) using *PP2AA3* (At1g13320) for normalisation. Each experiment was conducted using three biological replicates, each one of them including three technical replicates. Statistically significant differences were calculated using the tests described in the figure legend. Primer sequences are listed in Supplementary Table S1.

### MG132 and TUDCA treatments

7-days-old Arabidopsis seedlings from the *pHSFA1a:HSFA1a-GFP* lines (Col-0 and *hop1 hop2 hop3* mutant) that constitute the corresponding pairs were treated with 50 μg/ml MG132 or 0.5 mM TUDCA for 3 h. As controls, treatments with similar concentrations of DMSO were performed in parallel. After the treatments, the seedlings were collected and proteins were extracted in a buffer containing 50 mM Tris-HCl (pH 7.5), 150 mM NaCl, 5% (v/v) glycerol, 0.1% (v/v) Triton X-100, 0.5 mM phenyl-methanesulfonyl fluoride, and a protease inhibitor cocktail (Sigma). Proteins were quantified by the Bradford method, and equal amounts of protein were loaded in a gel and analyzed by western blot.

### Thermomorphogenesis assays

Analyses of hypocotyl elongation after the shift from control conditions to warm temperatures were performed as described in (Muñoz et al., 2022).

### RNA-Seq analyses

Three biological replicates of Arabidopsis seedlings from Col-0 and *hop1 hop2 hop3* mutant were grown under short-day conditions at 20°C for 4 days, side by side in a vertical position. After this initial growth phase, the plates containing the seedlings were divided in two groups. The first group was left at 20°C and the other one was transferred to 29°C for 24 h. Total RNA isolation was performed by the Trizol method (Ambion) followed by RNA cleanup by the RNeasy mini kit (Quiagen). Library construction (TruSeq Stranded mRNA) and sequencing (Illumina NovaSeq 6000) was performed at MACROGEN, resulting in 30-40 million paired-end reads (2 × 150) per sample.

Data analysis was performed on a Linux computer using a combination of command-line software tools, R packages from Bioconductor and Python scripts. FASTQ files were demultiplexed into individual libraries and total reads were mapped to the Arabidopsis genome (TAIR10) using the splice-aware read aligner HISAT2 (Kim et al., 2015), allowing only unique alignments and not more than two nucleotide mismatches. Minimum and maximum intron lengths were adjusted to 30 and 3,000 bp, respectively. The mapped RNA-seq reads were transformed into counts per transcript for each replicate/condition using feature counts (Liao et al., 2014), with the features annotated in the Arabidopsis GTF file (TAIR10.45). Differential expression analysis, based on the negative binomial distribution, were performed using DESeq2 R package (version 1.4.2).

## RESULTS

### The three members of the Arabidopsis HOP family interact in vivo with AtHSFA1a

It was previously reported that different members of the AtHSP90 family interact with AtHSFA1a AtHSFA1b and AtHSFA1d when expressed in epidermal onion cells and Arabidopsis protoplasts (Yoshida et al., 2011). In addition, it was described that treatment with the HSP90 inhibitor geldanamycin (GDA) inhibits the transcriptional activation under severe heat stress of the HSFA1-responsive gene *DREB2A*. Despite these data providing a preliminary link between HSP90 function and activity of the HSFA1 clade in the model plant, it remains unknown whether HSP90 is assisted by co-chaperones to carry out this function and, if so, which co-chaperone is involved. This latter is also a relevant question, since in eukaryotes HSP90 has a cohort of co-chaperones, some of them having a high selectivity to assist the folding of specific clients (Bose et al., 1996).

HOP is a co-chaperone of HSP90 that has a high selectivity for specific clients (Bhattacharya et al., 2020). As stated before, there are three HOP proteins in Arabidopsis, and it has been previously shown that the three members of the AtHOP family interact with HSP90 in plants (Fernandez-Bautista *et al*., 2018). However, apart from the Arabidopsis F-box protein TIR1, SNE and, probably, COI1 (Castellano, 2024; Mangano *et al*., 2023; Munoz *et al*., 2022; Muñoz et al., 2021), the identity of the other possible clients of HOP remains elusive in the plant kingdom. Therefore, in order to test if HSFA1a could be among the choice clients of HOP in Arabidopsis and to explore the selectivity of AtHOP members in this possible interaction, we expressed Flag-tagged versions of HOP1, HOP2 and HOP3, along with an HA-tagged version of HSFA1a in *N. benthamiana* leaves and we carried out co-immunoprecipitation analysis using anti-Flag beads. As observed in Figure 1A, the HSFA1a specifically co-immunoprecipitates with the three members of the Arabidopsis HOP family, suggesting that HOP1, HOP2 and HOP3 interact in planta with the HSFA1a.

**Figure 1.**
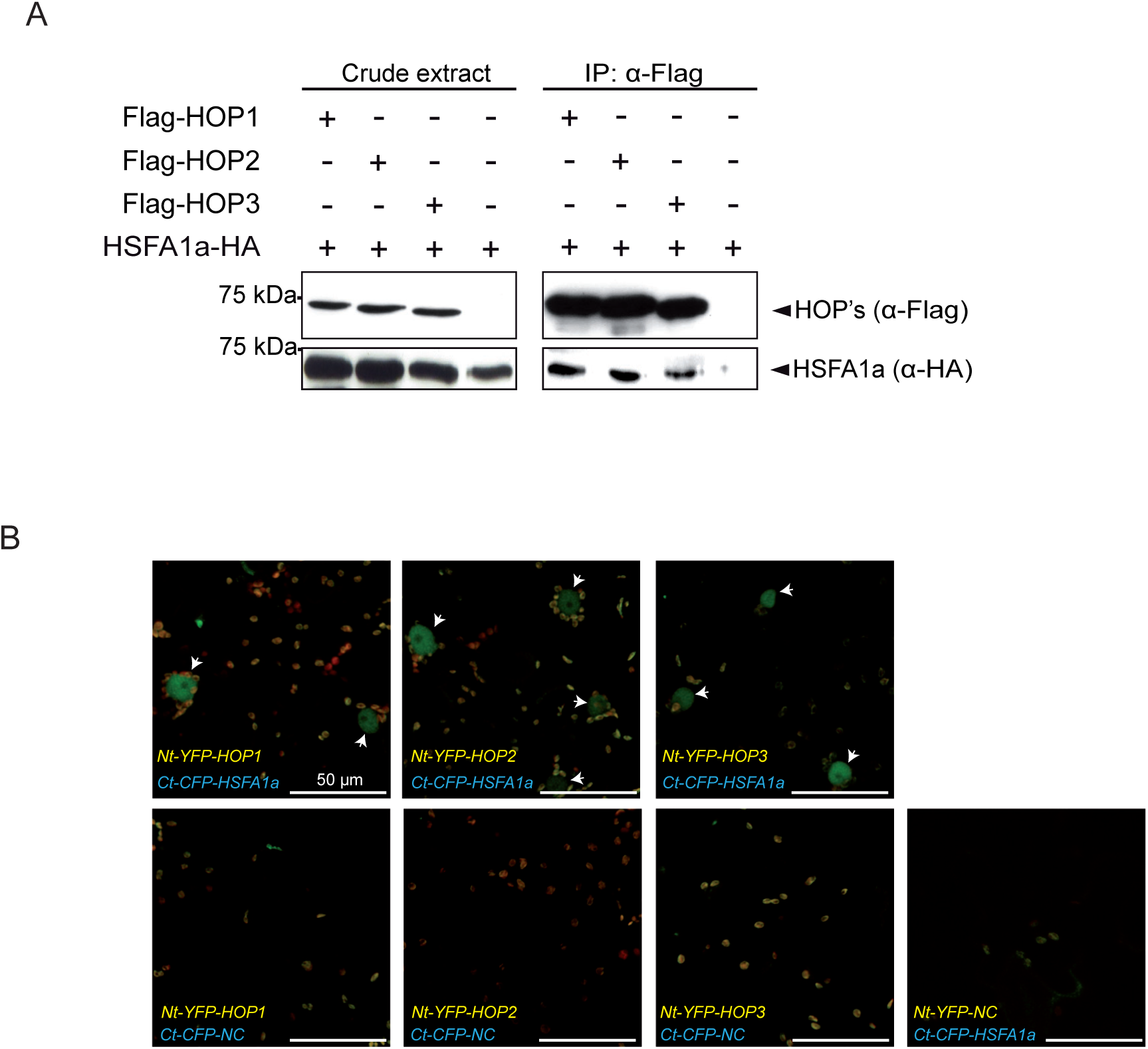
AtHSFA1a interacts with the three members of the Arabidopsis HOP family in vivo. (A) Co-immmunoprecipitation analyses. Protein extracts (crude extracts) from *N. benthamiana* leaves transiently expressing, under the control of the *35S* promoter, different combinations of Flag-AtHOPs and AtHSFA1a-HA tagged proteins were subjected to immunoprecipitation using anti-Flag beads. The presence of the different proteins in the crude extracts and in the eluted fractions from HOPś immunoprecipitations (IP:a-Flag) was analysed by western blot using anti-HA and anti-Flag antibodies. (B) Confocal images of BiFC analyses. Different combinations of AtHOPs fused to N-terminal of YFP and AtHSFA1a fused to C-terminal CFP fragments were expressed together or along with negative controls (NC) in *N. benthamiana* leaves. Reconstituted YFP-CFP fluorescence (detected as green fluorescence in the nuclei) is highlighted by arrows. Chloroplastś autofluorescence is observed as yellow small dots. In all cases results are representative of n=3 independent experiments.

### HOP proteins are not involved in the localization of the AtHSFA1a, but modulate its accumulation

The interaction of the HSFA1a with HOP proteins may suggest that HOP could somehow modulate some aspect of HSFA1a function or regulation. One of the most interesting aspects of HSF1 regulation in mammals is its retention in the cytoplasm in the absence of stress through the binding to HSP70 and HSP90 (Masser et al., 2020). Interestingly, previous reports have shown that the three members of the AtHOP family are mainly localized in the cytoplasm under control conditions, although a reduced localization in the nucleus is also observed (Fernandez-Bautista *et al*., 2018). This makes possible that the HOP cytoplasmic fraction could somehow help to retain the HSFA1a in the cytoplasm, impeding its transport to the nucleus in the absence of stress.

To test this hypothesis while exploring the HOPs/HSFA1a interaction by a different technique, we carried out bimolecular fluorescence complementation (BIFC) assays. For this, we expressed the proteins HOP1, HOP2 and HOP3 fused to the N-terminal part of YFP (Nt-YFP-HOPs) along with HSFA1a fused to the C-terminal part of CFP (Ct-CFP-HSFA1a) in *N. benthamiana* leaves

As observed in Figure 1B, when the fusion proteins of HOPs and HSFA1a are expressed together, a green signal, which denotes the physical interaction and complementation between the YFP and CFP moieties driven by the interaction of HOPs and HSFA1a, is observed in the nucleus of the transformed cells. This complementation is not observed when either HOPs or HSFA1a fusion proteins were co-transformed with the corresponding negative controls. Collectively, these data further support that the three members of the HOP family interact in vivo with the HSFA1a, and that this interaction takes place mainly in the nucleus of the cells.

HOP is a co-chaperone of HSP70 and HSP90 and, in mammals, these chaperones retain the HSFA1 in the cytoplasm in the absence of stress (Masser *et al*., 2020). Thus, the above described data may suggest a different mechanism of regulation from that described in mammals. Therefore, to explore by different means if HOP could somehow modulate the subcellular localization of the HSFA1a, we decided to analyze whether the HSFA1a is retained in the cytoplasm in the absence of HOP. For this, we transformed Arabidopsis Col-0 and *hop1 hop2 hop3* transgenic plants with a construct expressing, under the control of the *HSFA1a* promoter, the genomic sequence of the *HSFA1a* fused to *GFP*. Two independent pair of lines (pair #1 and pair #2) were selected for these and further analyses. The pair of lines #1 showed a global higher level of expression of the *HSFA1a-GFP* reporter than the pair of lines #2. (Figure 2A, qRT-PCR analyses), Nevertheless, within each pair of lines, similar levels of the *HSFA1a-GFP* transgene between the Col-0 and *hop1 hop2 hop3* triple mutant backgrounds are observed (Figure 2A), which gave us the opportunity to compare the effect of the lack of HOP independently in both lines.

**Figure 2.**
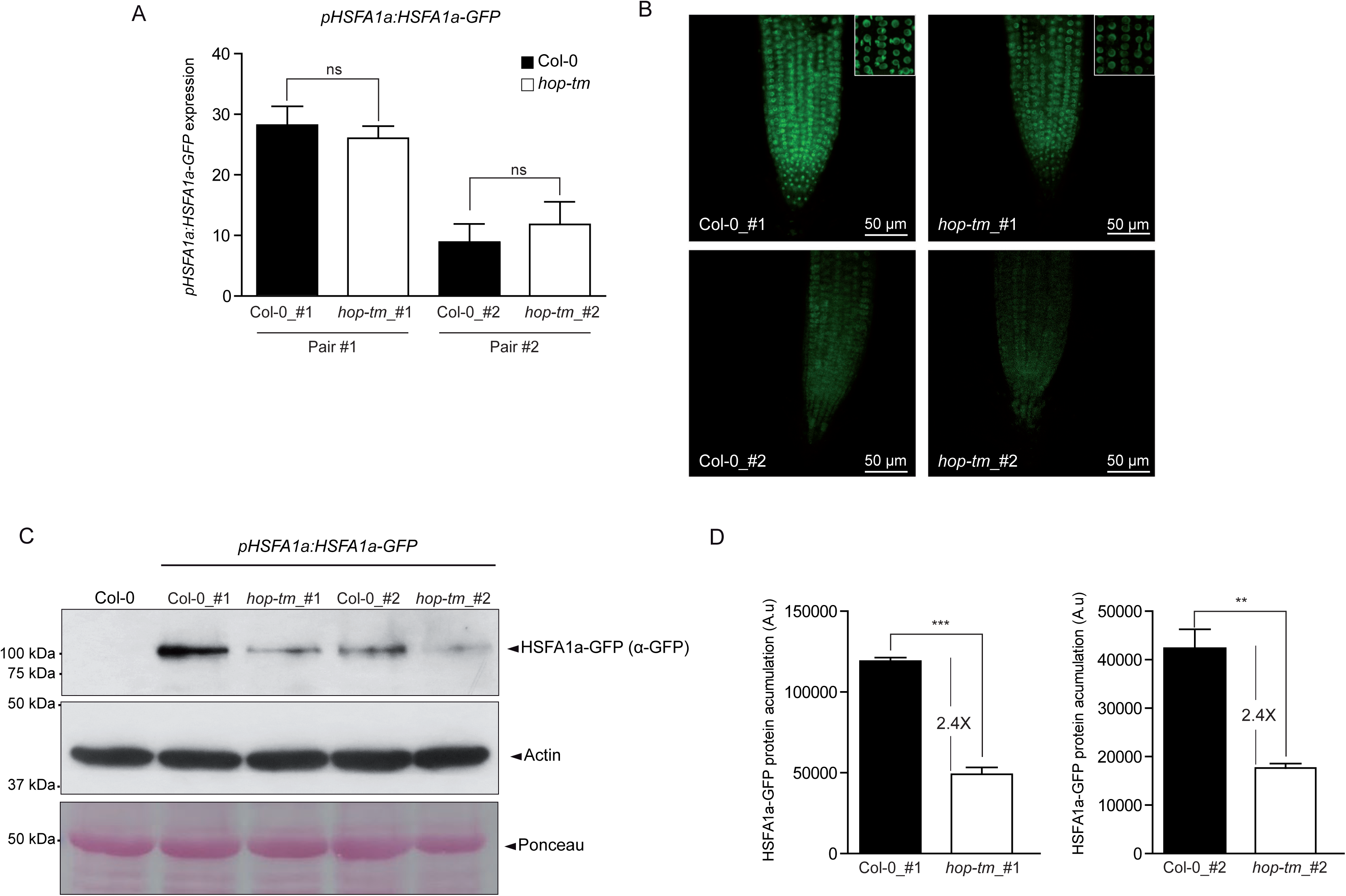
HOPs do not modulate the subcellular localization of the HSFA1a but do regulates HSFA1a stability under control conditions. (A) RT-qPCR analyses of the *HSFA1a-GFP* transgene in 7-day-old seedlings expressing the *pHSFa1a:HSFA1a-GFP* reporter construct in Col-0 or *hop1 hop2 hop3 (hop_tm)* backgrounds. Values are shown as mean±SD of n=3 independent experiments. Two different pair of lines (pair#1 and pair#2) displaying, within the pair, similar expression levels of the transgene *HSFA1a-GFP* in Col-0 and in the *hop1 hop2 hop3* mutant were selected for further analyses. (B) Representative confocal images of 7-day-old seedlings from the *pHSFA1a:HSFA1a-GFP* lines (Col-0 and *hop-tm*) that constitute the pairs#1 (upper panel) and pair#2 (lower panel). Close-up views from the images of the lines from pair #1 are also shown (right upper corner). Scale bars correspond to 50 µm. This experiment was done three times obtaining similar results. (C-D) Analysis of HSFA1a accumulation by western blot. Extracts from 7-day-old seedlings of the *pHSFA1a:HSFA1a-GFP* lines (Col-0 and *hop-tm*) that constitute the pair#1 and #2 were analysed by western blot using an anti-GFP antibody. Actin accumulation (analysed using an anti-ACTIN antibody) and Ponceau staining of the different extracts are provided as a loading control. Extract from Col-0 (non-expressing the reporter *pHSFA1a:HSFA1a-GFP*) is included as a negative control. (C) Representative results from the western blot analyses and (D) quantification (mean±SD values) from n=4 independent experiments as the one shown in (C). Statistically significant differences were calculated using (A) one-way ANOVA and the Tukey multiple comparison procedure or (D) Student’s t-test. ns, non-significant; *p < 0.05; **p < 0.01; ***p < 0.001).

To start this analysis, we analyzed the subcellular localization of the HSFA1a-GFP protein in Col-0 and in the *hop1 hop2 hop3* mutant using the pair of lines #1 and #2 in the absence of stress. Interestingly, and in contrast to the situation in mammals, where HSF1 shows a predominant cytoplasmic localization under control conditions, the HSFA1a-GFP fusion protein was localized in the nucleus at 20°C in the Col-0 background in both pair of lines (Figure 2B). No change in HSFA1a localization, which remains nuclear, was either observed in the *hop1 hop2 hop3* mutant background under such conditions (Figure 2B). The nuclear localization of the HSFA1a, and the co-localization of the HSFA1a with the bulk of HOP3 localized in the nucleus was further corroborated by co-expressing a different set of fusion proteins, in this case HSFA1a-Cherry and HOP3-GFP, in *N. benthamiana* leaves (Supplementary Figure 1). All these data reveal HSFA1a has a nuclear localization in the absence of stress and that, most probably, HOP proteins are not involved in the regulation of its subcellular localization.

Certainly, despite HSFA1a remaining nuclear, the accumulation of the HSFA1a-GFP protein seems to be drastically reduced in the *hop1 hop2 hop3* background in both pair of lines (lines #1 and 2) (Figure 2B). This reduced accumulation of the HSFA1a-GFP protein in the triple *hop* mutant was further corroborated by western blot (Figure 2C) and quantified (Figure 2D), being statistically significant in both pair of lines. Since the expression of the transgene in the wt and mutant background within each pair of lines is similar (Figure 2A), these data highly suggest that HOPs favor the accumulation of the HSFA1a under such conditions.

### HOP proteins module HSFA1a stability

Taking into account that HOP is a co-chaperone, it is possible that the reduced accumulation of the HSFA1a in the *hop1 hop2 hop3* mutant could be due to a defect in its folding that could be sensed by the quality control. If so, it is highly plausible that, in the *hop* triple mutant, the non-properly folded molecules of the HSFA1a would be targeted to their degradation by the proteasome. Therefore, to further test this hypothesis, we subjected the plants expressing HSFA1a-GFP in Col-0 and *hop1 hop2 hop3* mutant backgrounds to two different treatments: (1) with the proteasome inhibitor MG132, in order to test if the reduced accumulation of the HFA1a in the *hop* mutant is due to its degradation by the proteasome, and (2) with the synthetic chaperone TUDCA, to evaluate if this treatment could help to overcome the possible folding defect of HSFA1a, stabilizing the HSFA1a protein in the *hop1 hop2 hop3* mutant. As shown in Figure 3A, pair #1; Figure 3D, pair #2 and Figure 3C, quantification, no significant differences in the accumulation of HSFA1-GFP protein are observed in the wt background in the presence or absence of the MG132 treatment, suggesting that this protein is quite stable in Col-0 under control conditions. However, as described above, compared to the wt background, the HSFA1a-GFP protein is accumulated to lower levels in the *hop1 hop2 hop3* mutant (-MG132), reinforcing the idea that the HSFA1a turns unstable in the *hop1 hop2 hop3* mutant. Remarkably, this protein highly accumulates in the *hop* triple mutant plants (reaching similar levels to those of the wt plants) in the presence of MG132, indicating that, in the absence of HOP, the HSFA1a is partially degraded by the proteasome. Interestingly, as in the case of MG132, the treatment with the synthetic chaperone TUDCA prompts the accumulation of HSFA1a-GFP to wt levels in the *hop1 hop2 hop3* mutant (Figure 3D, pair #1; Figure 3E, pair #2 and Figure 3C, quantification). All these data highly indicate that HOP proteins are essential for the proper folding and maintenance of the stability of HSFA1a under control conditions, in such a way that, in the absence of these co-chaperones, the HSFA1a turns unstable and become degraded by the proteasome.

**Figure 3.**
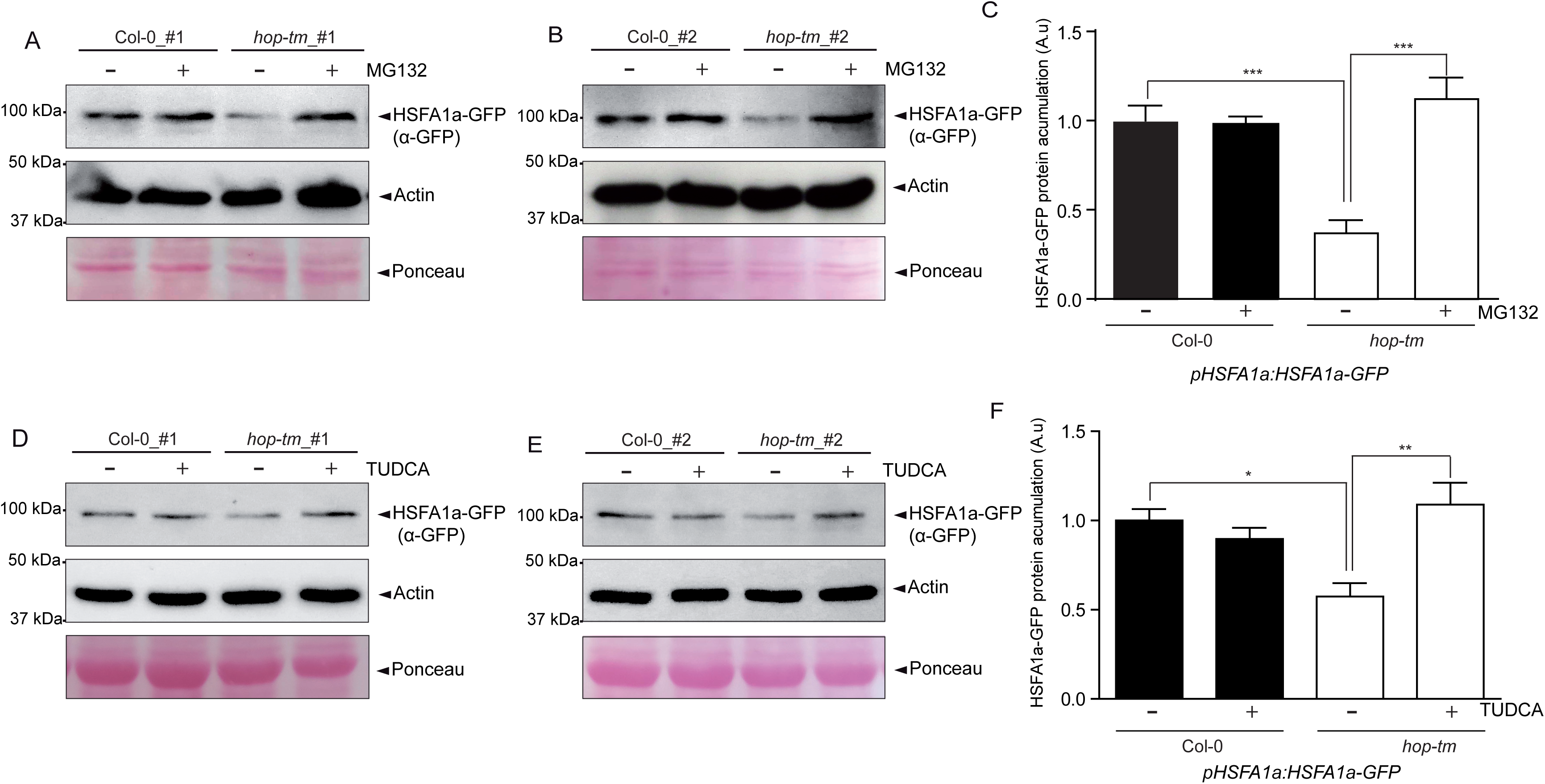
HOP is involved in the folding and stabilization of the HSFA1a. Analyses by western blot of HSFA1a accumulation in extracts from 7-day-old seedlings of the *pHSFA1a:HSFA1a-GFP* lines (Col-0 and *hop-tm*) that constitute the pair#1 and pair #2 using an anti-GFP antibody. Actin accumulation (analysed using an anti-ACTIN antibody) and Ponceau staining are provided as a loading control. (A-B) Representative western blot results and (D) quantification of n=3 independent experiments from the *pHSFA1a:HSFA1a-GFP* lines (Col-0 and *hop-tm*) of the pair#1 and pair#2 treated in the presence or absence of 50 μg/mL MG132 for 3 h. (D-E) Representative western blot results and (D) quantification of n=3 independent experiments from the *pHSFA1a:HSFA1a-GFP* lines (Col-0 and *hop-tm*) of the pair#1 and pair#2 treated in the presence or absence of 0.5 mM TUDCA for 3 h. In (C-F) values are expressed as mean±SEM. Statistically significant differences were calculated using one-way ANOVA and the Tukey multiple comparison procedure. *p < 0.05; **p < 0.01; ***p < 0.001.

### HOP proteins also play a role in the proper accumulation of the HSFA1a under warm temperature conditions

It is well established that the HSFA1a plays a main role in plant thermomorphogenesis (Cortijo *et al*., 2017). Therefore, in order to clarify the possible role of HOP in this process, we decided to analyze if HOP also influences the accumulation of HSFA1a under warm temperature or in the initial stages of the recovery from it. For this, we visualized and quantified GFP intensity as a measure of HSFA1a-GFP accumulation in roots from pair #1 that, as previously described, express to similar levels the construct *pHSFA1a:HSFA1a-GFP* in Col-0 and *hop1 hop2 hop3* mutant backgrounds. As observed in Figure 4A-B, according to the previous results, GFP intensity is significantly reduced in the roots of the *hop1 hop2 hop3* mutant compared to the wt at 20°C. This reduction in GFP intensity was also observed 3 h after the shift from 20°C to 29°C and once the challenge is ceased and plants are returned to control conditions. To corroborate these data, we carried out western blot analysis to directly analyze the accumulation of HSFA1a-GFP protein in seedlings. As shown in Figure 4C-D, similarly to the situation observed in roots, HSFA1a accumulation is also significantly reduced in the *hop1 hop2 hop3* seedlings under control conditions, during the initial stages of the challenge and during the recovery period. All these data suggest that HOP proteins are involved in the maintenance of the HSFA1a stability both at control and warm temperature conditions.

**Figure 4.**
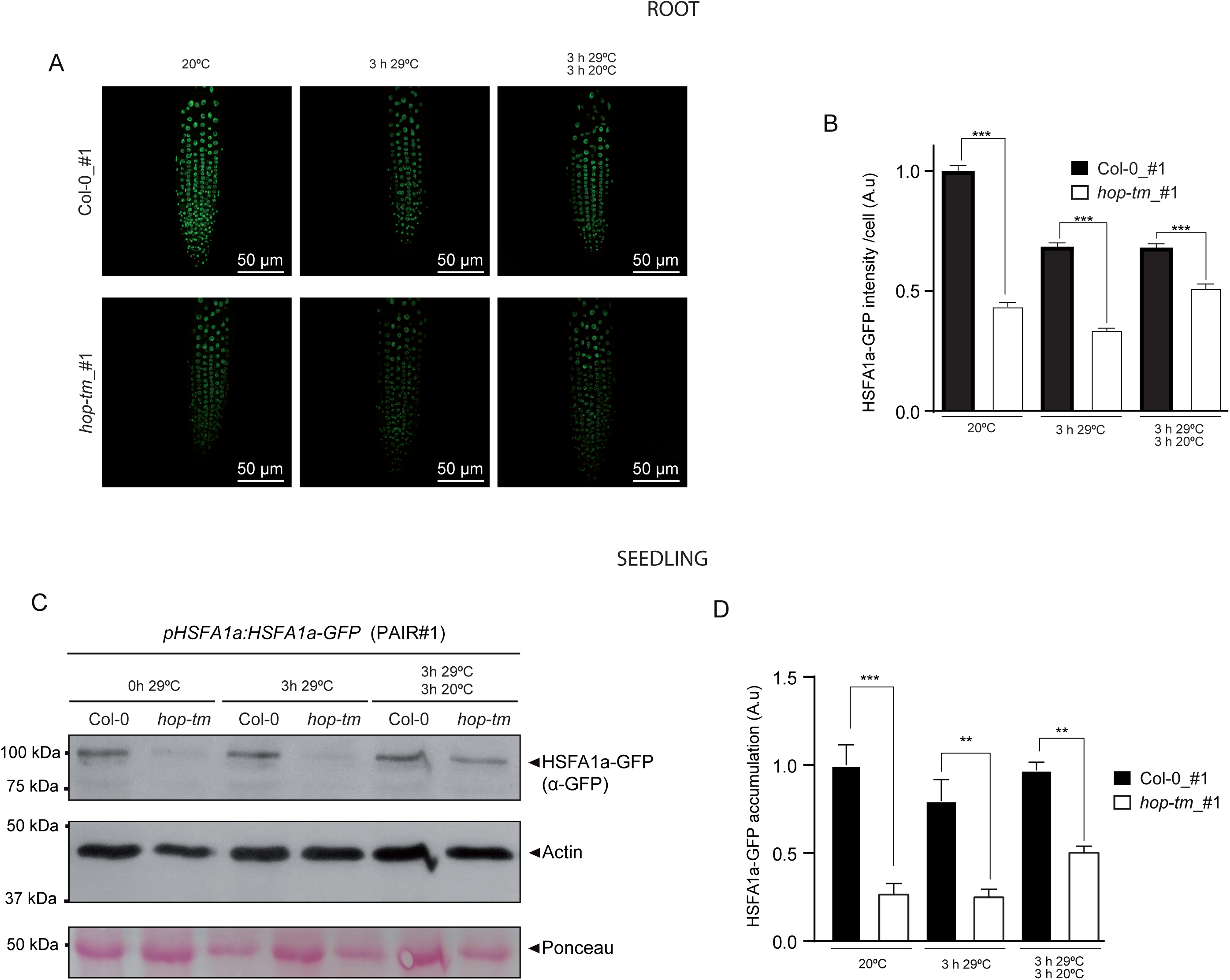
HOP modulates HSFA1a stability at warm temperatures and during recovery. (A) Analysis of HSFA1a accumulation by confocal microscopy in 7-day-old seedling from the *pHSFA1a:HSFA1a-GFP* lines (Col-0 and *hop-tm*) of pair#1, grown at 20°C, 3 h after the shift from 20°C to 29°C, and 3 h of growth at 20°C after the challenge (29°C). (A) Representative images and (B) quantification of GFP intensity per root nucleus n > 300 from the images and conditions described in (A). These experiments were performed three times obtaining similar results. (C-D) Analyses by western blot of HSFA1a accumulation in extracts from whole seedlings of *pHSFA1a:HSFA1a-GFP* lines (Col-0 and *hop-tm*) that constitute the pair#1 using an anti-GFP antibody, (a-GFP). Actin accumulation (analysed using an anti-ACTIN antibody) and Ponceau staining are provided as a loading control. (C) Representative western blot results and (D) quantification of n=3 independent experiments. In (B-D) values are expressed as mean±SEM. Statistically significant differences were calculated using (B) Kruskal-Wallis and the Dunn’s multiple comparison procedure and (D) one-way ANOVA and the Tukey multiple comparison procedure, (ns, non-significant; *p < 0.05; **p < 0.01; ***p < 0.001).

### HOP family is involved in hypocotyl elongation in response to warm temperatures and this phenotype is dependent on the proper establishment of the HSR

Our results indicate that HOP proteins bind to HSFA1a and modulate the folding and stabilization of the HSFA1a under control and mild heat stress conditions. Previous results from our group demonstrated that the *hop1 hop2 hop3* mutant shows defects in hypocotyl elongation during thermomorphogenesis (Mangano *et al*., 2023; Munoz *et al*., 2022); however, we do not know if the reduced accumulation of the HSFA1a could be determinant for the defect in hypocotyl elongation observed in the *hop1 hop2 hop3* mutant. Therefore, to assess this issue, we decided to analyze if hypocotyl elongation is altered under warm temperatures in the *hsfa1a* mutant, an aspect that to our knowledge has not been reported.

As shown in Figure 5A-B, no significant defects of hypocotyl elongation are observed under control conditions among the different genotypes (Col-0, and the mutants *hsfa1a* or *hop1 hop2 hop3*). However, both the *hsfa1a* and the *hop1 hop2 hop3* mutant show a reduced hypocotyl elongation under warm temperatures. These data corroborate that the reduced accumulation of the HSFA1a highly impinges on the thermomorphogenesis response.

**Figure 5.**
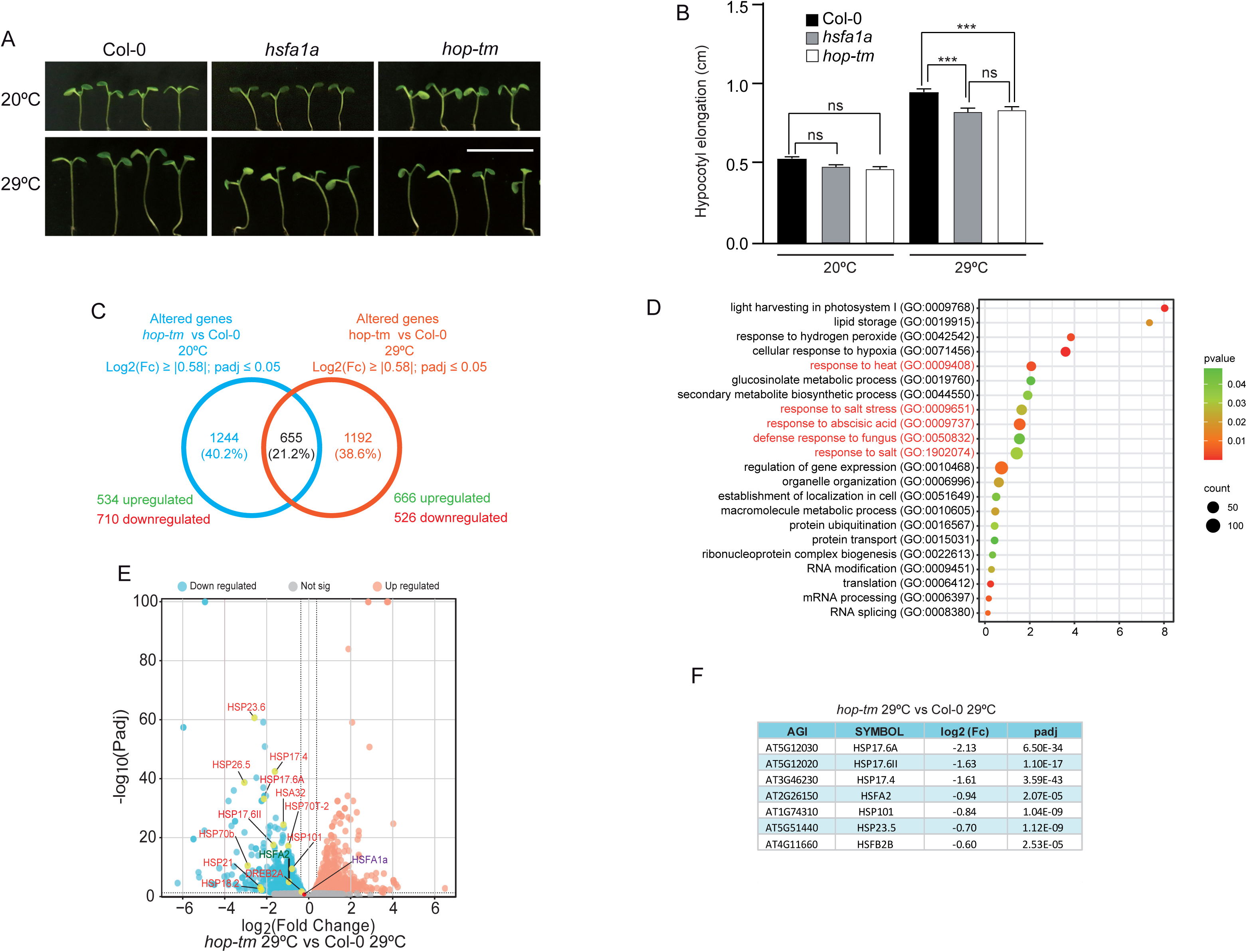
HOPs have a role in the onset of HSR and modulates the proper induction of the HSFA1a responsive genes. (A) Representative photographs and (B) quantification of hypocotyl elongation in seedlings from Col-0, the *hop1 hop2 hop3* triple mutant (*hop-tm*) and the *hsfa1a* mutant. Seedlings were grown for 4 days under short-day conditions at 20°C. After this time, seedlings were left to grow at 20°C or transferred to 29°C for additional 3 days. Data are expressed as mean±SEM from three independent replicates each containing *n* >50 seedlings for each genotype and condition. Statistically significant differences were calculated using one-way ANOVA and the Tukey multiple comparison procedure (ns, non-significant; **p < 0.01; ***p < 0.001). (C) Venn diagrams showing the number of overlapping and unique genes that are significantly changed (log2 (Fc) 2’ 10.581; padj :: 0.05) in the *hop1 hop2 hop3 (hop_tm)* mutant compared to Col-0 at 20°C or after their transfer to 29°C for 24 h. The number of upregulated genes (log2 (Fc) 2’ 0.58; padj :: 0.05) and downregulated genes (log2 (Fc) :: -0.58; padj :: 0.05) in in *hop-tm* compared to Col-0 at each temperature is also indicated. (D) Bubble plot showing the GO analysis of the significant altered genes in *hop-tm* compared to Col-0 at 29°C (log2 (Fc) 2’ 10.581; padj :: 0.05). This analysis was done using Fisher statistical test and FDR for the analysis of p-value. Colour code is shown at the right of the panel. Stress categories are highlighted in red. (E) Volcano plots showing the change in expression of the genes in *hop-tm* compared to Col-0 after the transfer to 29°C. Peach coloured dots represent significantly upregulated (log2 (Fc) 2’ 0.58; padj :: 0.05) and blue dots are downregulated genes (log2 (Fc) :: -0.58; padj :: 0.05). Some genes previously associated to the HSR are highlighted in yellow and specifically the HSFA2a is highlighted in green. The dot corresponding to HSFA1a is shown in red. Grey dots denote genes with a non-significant change. (F) Fold change and p-value of different known HSFA1a targets showing an altered expression in *hop-tm* compared to Col-0 when transferred at 29°C.

### HSR is highly altered in response to warm temperatures in the *hop1 hop2 hop3* mutant

It is well known that the HSFA1 clade, and specifically the HSFA1a, plays a major role in the transcription of multiple HSR responsive genes during thermomorphogenesis (Cortijo *et al*., 2017). Based on this, we hypothesized that, if the defects in thermomorphogenesis of the *hop1 hop2 hop3* mutant are associated to the reduced accumulation of the HSFA1a, the expression of the HSR will be altered in the *hop* triple mutant at warm temperatures. Thus, we decided to test if the alteration in the levels of the HSFA1a protein observed in the *hop1 hop2 hop3* mutant background could have an impact in the proper establishment of the HSR at warm temperature. For this, we performed RNAseq analysis on RNA obtained from seedlings of Col-0 and *hop1 hop2 hop3* mutant grown at 20°C or shifted to 29°C for 24 h. These analyses revealed that 1899 genes are differentially expressed (log2 (Fc) 2’ 10.581; padj :: 0.05) in the *hop1 hop2 hop3* mutant compared to Col-0 under control conditions (20°C), while 1847 genes are differentially expressed (log2 (Fc) 2’ 10.581; padj ::0.05) in the *hop1 hop2 hop3* mutant at warm temperature (29°C) (Figure 4C). Despite the number of altered genes in the mutant in both conditions being quite similar, we observed that the identity of the genes and the associated functions to the altered genes in the *hop* mutant are largely distinct under control and mild-heat stress (Figure 4C and Supplementary Figures 2-3).

From the 1192 differentially expressed genes in the *hop1 hop2 hop3* at 29°C, 666 genes were up regulated (log2 (Fc) 2’0.58; padj :: 0.05) while 526 genes displayed a reduced expression in the *hop1 hop2 hop3* mutant at warm temperature (log2 (Fc) :: -0.58; padj :: 0.05) (Figure 4C). Gene ontology (GO) analyses demonstrated that these sets are highly enriched in GO categories related to stress, including response to heat (GO:0009408) (Figure 5D). Interestingly, among the genes whose expression is significantly reduced in the *hop1 hop2 hop3* mutant under warm temperature, different genes associated to the HSR, such as *HSFA2*, *HSP101*, *HSPl7.4*, *HSP18.2*, *HSA32* or *DREB2A*, are clearly identified (Figure 4E). All these data demonstrate that HOP is required for the proper onset of the HSR in response to warm temperature Intriguingly, some of the genes which are altered in the *hop1 hop2 hop3* mutant at 29°C, such as *HSFA2* and *HSP101*, are well-known targets of the HSFA1a. This prompted us to analyze in more detail the link between HOP and HSFA1a activity. Previous ChiP-seq analyses of HSFA1a, allowed the identification 1371 direct HSFA1a target genes during thermomorphogenesis (Cortijo *et al*., 2017). Using this information, we analyze the presence of those HSFA1a-responsive genes among the altered *hop1 hop2 hop3* compared to Col-0 at 29°C. Interestingly, the expression of 103 HSFA1a-regulated genes is significantly altered in the *hop1 hop2 hop3* mutant at 29°C (log2 (Fc) 2’ 10.581; padj :: 0.05) (Supplementary Table 2), indicating that HOP activity modulates HSFA1a function under mild-heat stress. Some of the better known HSFA1a targets, along with their fold change in the *hop1 hop2 hop3* mutant compared to Col-0 at 29°C, are displayed in Figure 5F. Noteworthy, no significant change in the expression of the *HSFA1a* gene is observed between the Col-0 and the *hop1 hop2 hop3* mutant either at 20°C (log2 (FC) = 0.096; padj = 0.896) or 29°C (log2 (FC) = -0.04; padj = 0.938). These data, along with the reduced accumulation of HSFA1a protein in the *hop1 hop2 hop3* mutant, highly suggest that the misexpression of the HSFA1a-dependent genes observed in the *hop1 hop2 hop3* mutant under warm temperature is not due to a down-regulation of the *HSFA1a* gene expression but, instead, could be due to a decreased accumulation of the HSFA1A protein. Moreover, these results provide evidence of the relevance of HOP function in the stabilization of the HSFA1a protein and in the proper onset of the HSR in response to warm temperature.

## DISCUSSION

### AtHOP does not modulate the HSFA1a localization in response to mild temperature

HOP is a co-chaperone of HSP70 and HSP90 (Castellano, 2024; Johnson *et al*., 1998; Odunuga *et al*., 2004; Toribio *et al*., 2020); therefore, it is highly probable that its function is associated to the activity of these chaperones. HSP70 and HSP90 have been involved in different aspects of proteostasis stresses, and one of these aspects, following the canonical titration by chaperone model, includes the control of the subcellular localization of the HSF1 (Masser *et al*., 2020). Indeed, it has been proposed that, in mammals, the HSF1 is retained in the cytoplasm through the binding to HSP70 and HSP90 in the absence of stress. However, proteostasis stresses, such as heat stress, usually course with a high accumulation of misfolded proteins, which compete with the HSF1 for chaperone binding. Under such conditions, the high accumulation of misfolded proteins force the chaperones to release the HSF1, which now accumulates in the nucleus and binds the promoters of its responsive genes, allowing the onset of the HSR (Anckar and Sistonen, 2011; Masser *et al*., 2020; Voellmy and Boellmann, 2007). Remarkably, it has been recently reported that the nuclear accumulation of HSF1 is reduced by depletion of HOP in mammalian cells, which indicates that, in addition to the HSP70 and the HSP90 chaperones, the co-chaperone HOP could be also involved in the tritation of HSF1 in these specific eukaryotes (Chakraborty and Edkins, 2020).

As in the case of the mHSF1, it has been previously reported that the only member of the HSFA1 family in tomato increases its nuclear localization in response to heat (Scharf et al., 1998). Similar results were also obtained for the HSFA1b, HSFA1c and HSFA1e in Arabidopsis, being this accumulation dependent on HSP90 (Yoshida *et al*., 2011). Interestingly, the subcellular localization of AtHOP proteins also show a similar subcellular localization to mHSF1, since AtHOPs have a predominant cytosolic localization compared to the nucleus under control conditions, but highly accumulate in the nucleus under severe heat (Fernandez-Bautista *et al*., 2018). These parallelisms open the possibility that, if the AtHSFA1a is regulated in a similar way as mHSF1, HOP could somehow participate in the subcellular localization of HSFA1a in response to heat in Arabidopsis.

Although this theory could not be ruled out in the case of the AHSFA1 members b, d and e, this does not seem to be the case for the AtHSFA1a. Our experiments carried out in Arabidopsis and *N. benthamiana* show that the bulk of AtHSFA1a is already localized in the nucleus in the absence of stress. Remarkably, these observations are in agreement with the nuclear localization of the AtHSFA1a during its ectopic expression under a constitutive promoter in onion epidermal cells (Yoshida *et al*., 2011). Furthermore, in contrast to the dependence on HSP90 for the transport of

AtHSFA1b, AtHSFA1d and AtHSFA1e to the nucleus, AtHSFA1a localization is independent of HSP90 or HOP. Certainly, AtHSFA1a remains nuclear in the presence of GDA and in the *hop1 hop2 hop3* mutant ((Yoshida *et al*., 2011), Supplementary Figure 4). All together our data suggest that, although it is plausible that HSFA1b, HSFA1d and AtHSFA1e follow the canonical titration by chaperone model in Arabidopsis, AtHSFA1a does not. These differences make highly interesting to analyze why the HSFA1a interacts with the co-chaperone HOP and if this interaction has an effect on HSFA1a regulation.

### Role of HOP in HSFA1a stabilization

Our data indicate that HSFA1a could be considered a client of HOP in Arabidopsis. This statement is based on different observations: (1) The three members of the HOP family bind to the HSFA1a. (2) As revealed by the increased accumulation of the HSFA1a in the *hop1 hop2 hop3* mutant in the presence of MG132, the HSFA1a becomes unstable and degraded by the proteasome in the absence of HOP. (3) The reduced accumulation of AtHSFA1a in the *hop1hop2 hop3* mutant is reverted in the presence of the synthetic chaperone TUDCA, suggesting that the role of HOP in this process is related to the folding of the HSFA1a. Altogether these data indicate that HOP intrinsically promotes the folding of the HSFA1a, which allows this transcription factor to become more stable.

Interestingly, our experiments suggest that HOP is required for the stabilization of the HSFA1a under control conditions and also during the stress, indicating that HOP-dependent stabilization is intrinsically needed to maintain the levels of the HSFA1a in plant cells. Although our data reflect that HOP proteins participate in the folding of the HSFA1a, we still do not know how HOP affects the structure of this transcription factor. It has been reported that, in mammals, the stability of the HSF1 is regulated by different post-transcriptional modifications (PTMs) (Dayalan Naidu and Dinkova-Kostova, 2017). In this sense, it has been described that the phosphorylation of the residues S303 and S307 marks HSFA1 to degradation by the SCF/FBXW7a complex in heat stressed late passage and serum –free human fibroblasts (Bonelli et al., 2001). Moreover, the phosphorylation of the HSF1 at S216 has been associated to its degradation by the SCF/*β*-TrCP during mitosis. Interestingly, HSFA1 degradation is reduced by acetylation at its K208 and K298 residues by the acetyltransferase CREB-binding protein/p300-type histone acetyltransferase (EP300) (Raychaudhuri et al., 2014). Based on the relevance of PTMs in HSFA1 stability in mammals, it is possible to speculate that plant HOPs could propitiate an intermediate folding stage in which the residues involved in HSFA1a degradation are hindered, or, instead, in which the sites involved in the HSFA1a stability are exposed. In this sense, it is interesting to point out that *hac1-2*, an Arabidopsis mutant in a gene that encodes a p300-type histone acetyltransferase (a protein that promotes HSF1 stability in mammals), shows a reduced expression in response to heat of two important HSFA1a target genes (*HSFA2* and *HSP70*). This reflects that this mutant shows a defect in the establishment of the HSR and, in accordance, this mutant shows a reduced short-term acquired thermotolerance (Sharma et al., 2021). At this stage, it is difficult to know whether, in Arabidopsis, the folding by HOP promotes HSFA1a stability through the binding and acetylation by HAC1. Indeed, knowledge on HSFA1 regulation by PTMs is extremely scarce in plants and no events of PTM by acetylation have been described regarding AtHSFA1a. Nevertheless, this is an interesting link that seems worth to study in the future. Upcoming experiments could also provide evidence on the role of these and other PTMs in HSFA1a regulation.

Different members of the Arabidopsis HOP family show a certain degree of specialization for specific functions. This is the case of HOP3, since it seems to be the only member involved in plant defense against *T. urticae* infestation so far. Furthermore, a prevalent role of HOP2 and HOP3 (over HOP1) has been described in GA signaling (Mangano *et al*., 2023; Muñoz *et al*., 2021). In this work, we observe that the three members of the AtHOP family interact in vivo with the HSFA1a. In addition, we have previously described that HOP1, HOP2 and HOP3 participate in hypocotyl elongation in response to mild temperatures (Mangano *et al*., 2023; Munoz *et al*., 2022). These data allow us to speculate that the three members of the Arabidopsis HOP family contribute to maintain the HSFA1a stability at warm temperature. Information about the regulation of the members of the AtHOP family is extremely scarce; however, there are already some clues that suggest that they may be differentially regulated (Castellano, 2024). For this reason, although we can speculate that the three members could participate in the response, we still do not know if some members could play a prevalent or selective role over the others.

### Impact of HOP-dependent HSFA1a stabilization on HSR

Our results indicate that HOP-dependent stabilization of the HSFA1a is not enough to induce the HSR under control conditions. Indeed, a challenge (in this case an increase of temperature from 20 to 29°C) is needed to observe the impact of HOP activity on the HSR. This seems to suggest that although HOPs already stabilize the HSFA1a under control conditions, HOPs do not render the HSFA1a completely functional, and this transcription factor needs to be further activated to promote the response.

Previous reports indicate that the HSR is induced in Arabidopsis seedlings by heat, even when the seedlings have been treated before the challenge with cycloheximide (an inhibitor of translation) (Yamada et al., 2007). These results highly suggest that the activation of the HSR depends on transcription factors (most probably, on the HSFA1 family) that are already present under control conditions (Yamada *et al*., 2007). Our results show that HSFA1a is already stabilized by HOP in the absence of stress, and that this contributes to the proper onset of the HSR under mild temperatures. In this sense, it is possible to speculate that the reduced levels of the HSFA1a under control conditions observed in the *hop1 hop2 hop3* triple mutant probably hampers the initial re-programming of the HSR during the response to heat in the mutant lines

### Role of HSP90 and HOP in the onset of the HSR

In Arabidopsis, HSP90 inhibition with GDA and radicicol actives the HSR, inducing more than 100 genes related to the HSR in the absence of heat shock (Yamada *et al*., 2007). Moreover, these inhibitors, most probably through the activation of the HSR, promote heat tolerance to high temperatures in Arabidopsis seedlings (Yamada *et al*., 2007; Yamada and Nishimura, 2008). Similar results were obtained with a different HSP90 inhibitor, Monocillin I (MON). Certainly, the application of MON leads the expression of major components of the HSR, such as the genes *HSP101* and *HSP70*, and promotes heat tolerance of Arabidopsis seedlings. In addition, co-cultivation of the MON-producing fungus *P. quadriseptata* with Arabidopsis leads to enhanced heat tolerance of Arabidopsis (McLellan et al., 2007). Taken together, these data demonstrate that, similarly to the situation in other eukaryotes, the HSP90 plays an overall inhibitory role in HSR activation in Arabidopsis.

It was previously showed that HOP1, HOP2 and HOP3 interact with HSP90 (Fernandez-Bautista *et al*., 2018). Despite HOPs being considered important HSP90 co-chaperones, the involvement of HSP90 in HSFA1a stability does not make trivial the participation of HOP in this process. Certainly, recent investigations in mammals and yeast revealed that, far from sharing the full network of HSP90 clients, HOP seems to be specialized in the folding of an extremely selective subgroup of them (Bhattacharya and Picard, 2021). In addition, it has been described that HOP could play HSP90-independent roles, which also highlights the relevance of this association (Gebauer et al., 1998; Odunuga *et al*., 2004).

In this article we observe a reduction in the HSFA1a levels when plants are treated with the HSP90 inhibitor GDA (Supplementary Figure 4), suggesting that HOP, along with HSP90, participates in maintaining the stability of the HSFA1a (Supplementary Figure 4). For this reason, it is intriguing how HOP could be needed for the efficient set up of the HSR while, in contrast, HSP90 plays an opposite role in the same process. This aspect becomes even more fascinating, taking into account that HSP90 and HOP bind both to HSFA1a in vivo (Yoshida *et al*., 2011). Having in mind that HOP is a co-chaperone of HSP90 and HSFA1a is a master regulator of the HSR associated to thermomorphogenesis (Cortijo *et al*., 2017), it is possible to speculate that HSP90 is involved in two sequential steps of HSFA1a regulation (Figure 6): a first step in which the HSP90 and HOP would be required in promoting the initial folding state of the HSFA1a. This folding would be required for its stabilization (turning the HSFA1a unstable in the absence of HOP). In addition, HSP90 would also be involved in a second step of the HSFA1A regulation, inhibiting the activity of this factor until the heat is sensed. This second step would be independent of HOP, since we do not observe, in contrast to which happens by the inhibition of the HSP90 with GDA, the activation of the HSR in the absence of stress in the *hop1 hop2 hop3* mutant. In this model, it is possible that HSP90 and HOP only exert their folding function in a reduced proportion of the HSFA1a molecules. In this regard, it is important to note that although we observe a decreased accumulation of the HSFA1a in the *hop1 hop2 hop3* mutant, this mutant does not exhibit a null accumulation of this transcription factor. Instead, the function of the HSP90 in the second step would affect the general bulk of the HSFA1a. This will make this second step more relevant in the general output of the response, which may explain the general inhibitory effect of HSP90 on the HSR.

**Figure 6.**
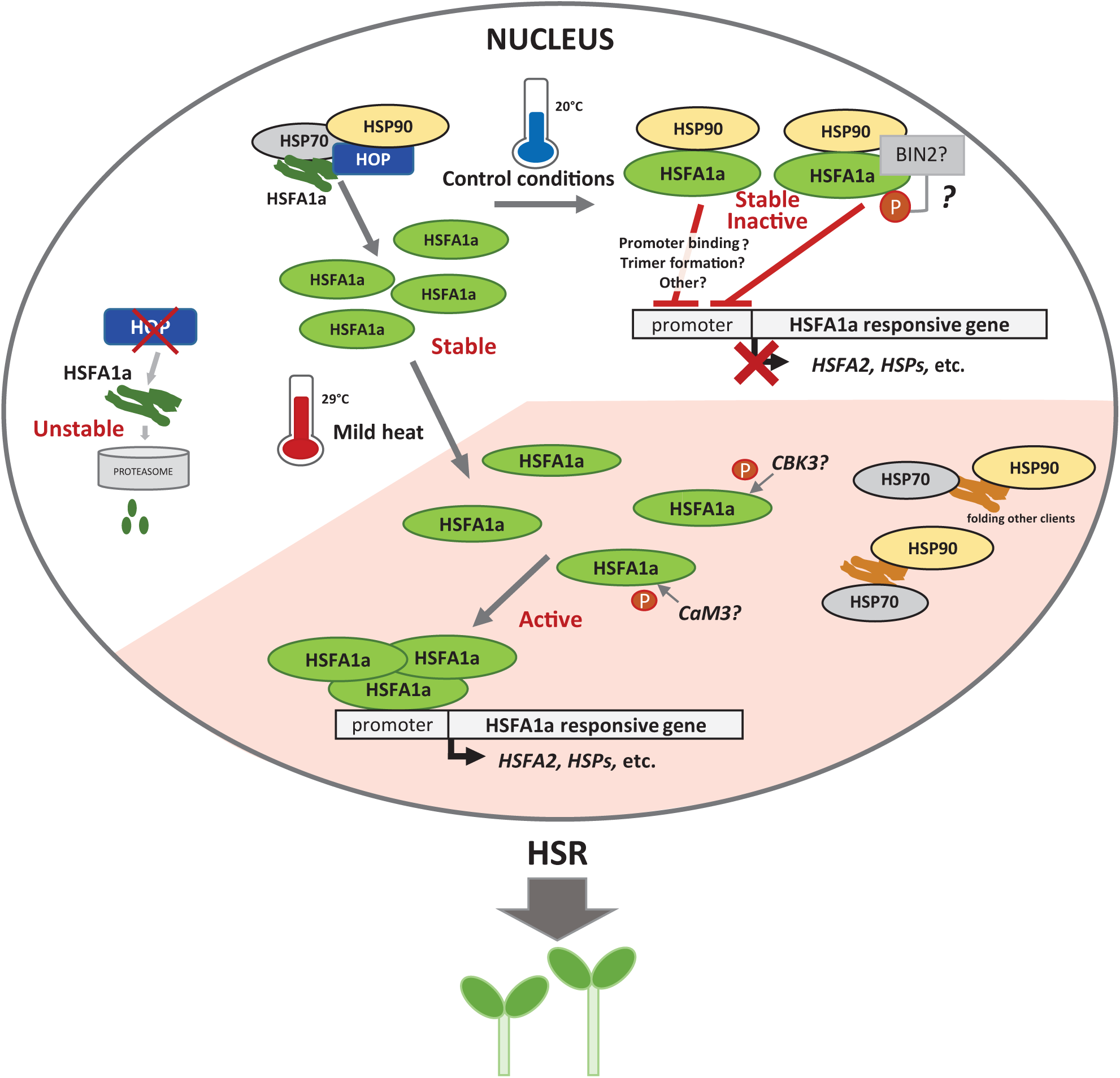
Proposed model of the roles of HOP and HSP90 in the activation of the HSFA1a in Arabidopsis. AtHOPs localize in the cytoplasm and the nucleus under control conditions. In the nucleus HOPs interact with HSFA1a (most probably generating a complex with HSP70 and HSP90, since it has been already described that HSP90 interacts with HSFA1a (Yoshida *et al*., 2011)). This complex favours the folding of the HSFA1a, stabilizing the HSFA1a and preventing its degradation by the proteasome. By promoting HSFA1a stability, HOPs contribute to the accumulation of the HSFA1a under control conditions and under stress. However, the accumulation of HSFA1s is not enough to promote the activation of the HSR in the absence of stress, since, as revealed by treatments with HSP90 inhibitors (Yamada *et al*., 2007), the HSR remains blocked by HSP90 activity under control conditions. It is highly possible that, as described in mammals, the inhibitory effect of the HSP90 over the HSFA1a is released under stress as the HSP90 is then recruited to maintain the general homeostasis. Under such conditions, the HSFA1a binds to the promoters of the HSFA1a responsive genes, triggering the HSR. How HSP90 inhibits the activity of the HSFA1a is already unknown; however, it has been reported that the kinase BIN2, a known client of HSP90, binds to all members of the AtHSFA1 family and phosphorylates the AtHSFA1d at S56, inhibiting the binding of this transcription factor to the promoter of *HSP18.2* (a known HSR responsive gen)(Luo et al., 2022). Interestingly, this residue is also conserved in the AtHSFA1a, which opens the possibility that this kinase, through its binding to HSP90, could be inhibiting the HSFA1a activity under control conditions. It is also plausible that HSFA1a activation requires phosphorylation events whose residues could be hindered by its association with HSP90. Interestingly, two kinases, the Calmodulin (CaM)-binding protein kinase CBK3 and CaM3, have been related to the HSFA1 family and proposed positive regulators of the HSR in plants (Liu et al., 2008; Zhang et al., 2009).

It is relevant to establish that the experiments with the tagged HSFA1a allow us to set the focus on the HSFA1a, an important actor of the thermomorphogenesis response in Arabidopsis. Noteworthily, AtHSFA1a has multiple peculiarities compared with the other members of the HSFA1 family. Indeed, the HSFA1a shows a constitutive nuclear localization that does not depend on the HSP90 activity. In contrast, the HSFA1b, HSFA1d and HSFA1e are localized in the cytoplasm or in the cytoplasm and nucleus under control conditions, but increase their accumulation in the nucleus under heat stress. Moreover, the retention of these latter members in the cytoplasm is dependent on HSP90, since the treatment with GDA promotes their accumulation in the nucleus in the absence of stress (Yoshida *et al*., 2011). It has been described that the HSFA1a and the HSFA1d play prevalent roles over the rest of the members in response to heat shock (severe heat) (Nishizawa-Yokoi et al., 2011); however, as far as we know, the relevance of AtHSFA1b, AtHSFA1d and AtHSFA1e in response to mild-heat stresses has not been specifically addressed. If some of these members are relevant, further studies will be needed to understand whether HOP could also contribute to their folding and stability. Furthermore, taking into account that HOPs, as these specific AtHSFA1 factors, increase their nuclear accumulation in response to high temperatures, it would be interesting to analyze if HOP could also modulate this aspect of their regulation.

Along with the HSFA1a, the F box proteins TIR1 and SNE involved in the auxin and GA pathway have been proposed as strict clients of HOP in Arabidopsis (Mangano *et al*., 2023; Munoz *et al*., 2022). In addition, it has been recently described that AtHOP1 and AtHOP2 bind to BIN2 and modulate BIN2 nuclear localization in response to BRs, which points out to the involvement of HOP in the BRs pathway (Zhang et al., 2022). Interestingly, all these pathways, along with PIF4, coordinately influence the thermomorphogenesis output (Quint *et al*., 2016). This makes possible that the phenotype observed in the *hop* mutants is derived from their global effect on these routes. In our assays we clearly observe that the HSR is altered under mild heat stress conditions in the *hop1 hop2 hop3* mutant. This along the binding of HOP to HSFA1a and its degradation in the *hop1 hop2 hop3* mutant highly suggest that HOP plays a major role over this specific route in this process. Whether HOP could affect the folding of other important signaling proteins during thermomorphogenesis is still under investigation.

## ACKNOWLEDGEMENTS

We thank Alfonso Muñoz for critical reading of the manuscript. This work is supported by the projects PID2021-126956OB-I00 from MICIU, by “Severo Ochoa Programme for Centres of Excellence in R&D” from the Agencia Estatal de Investigación of Spain (grants SEV-2016-0672 and CEX2020-000999-S to the CBGP). In the frame of this latter program R. T. was supported with postdoctoral contracts. In addition, A. N was funded by Comunidad Autónoma de Madrid (CAM) with a contract from “Programa de Empleo Juvenil” PEJ-202-AI/BIO-21922.

---

**Supplementary Figure 1.**
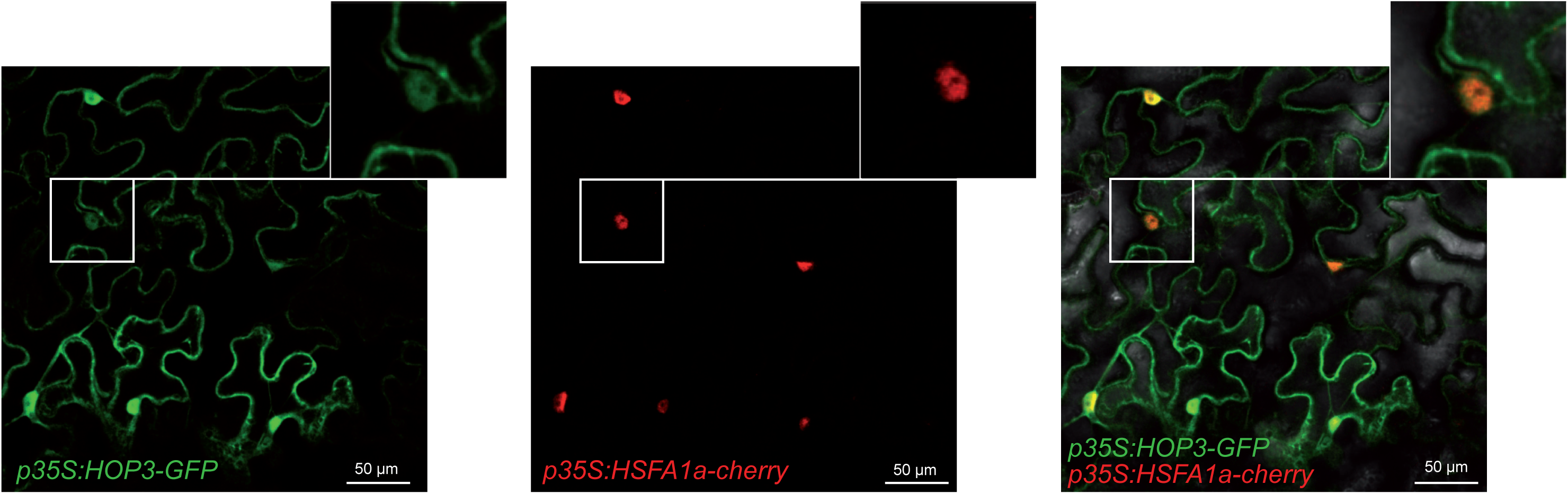
HSFA1a co-localizes with HOP3 in the nucleus under control conditions. Subcellular co-localization analyses of different combinations of the tagged AtHOP3- GFP and AtHSFA1a-CHERRY expressed in *N. benthamiana* leaves. Scale bars correspond to 50 µm. Close-up views of the nuclei are shown in the right corner of each figure. These experiments were performed three times obtaining similar results.

**Supplementary Figure 2.**
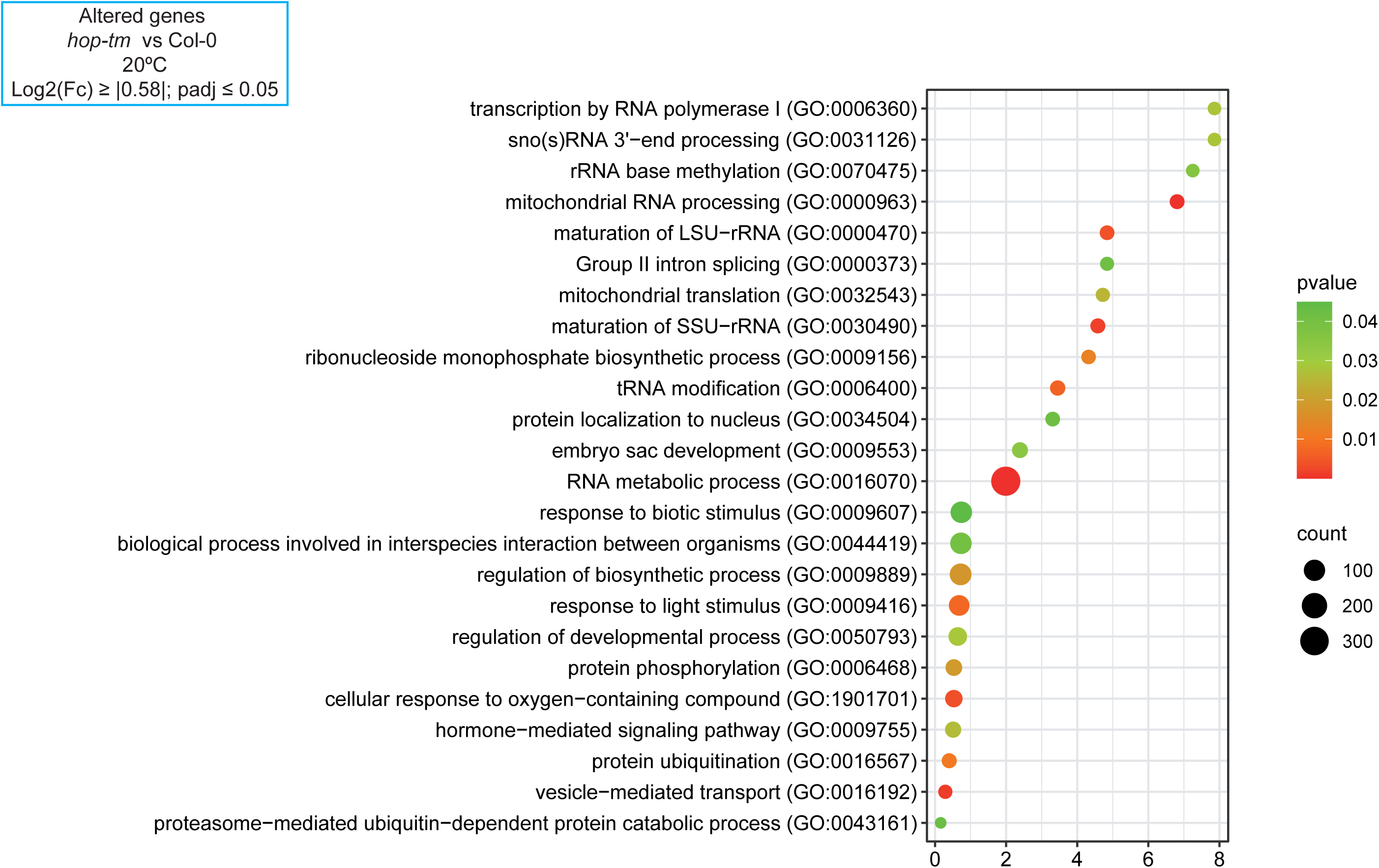
GO enrichment of the genes altered in the *hop1 hop2 hop3* mutant at control conditions. Bubble plot showing the GO analysis of the significant altered genes in *hop1 hop2 hop3* triple mutant (*hop-tm)* compared to Col-0 at 20°C (log2 (Fc) 2’ 10.581; padj :: 0.05). This analysis was done using Fisher statistical test and FDR for the analysis of p-value. Colour and count codes are shown at the right of the panel.

**Supplementary Figure 3.**
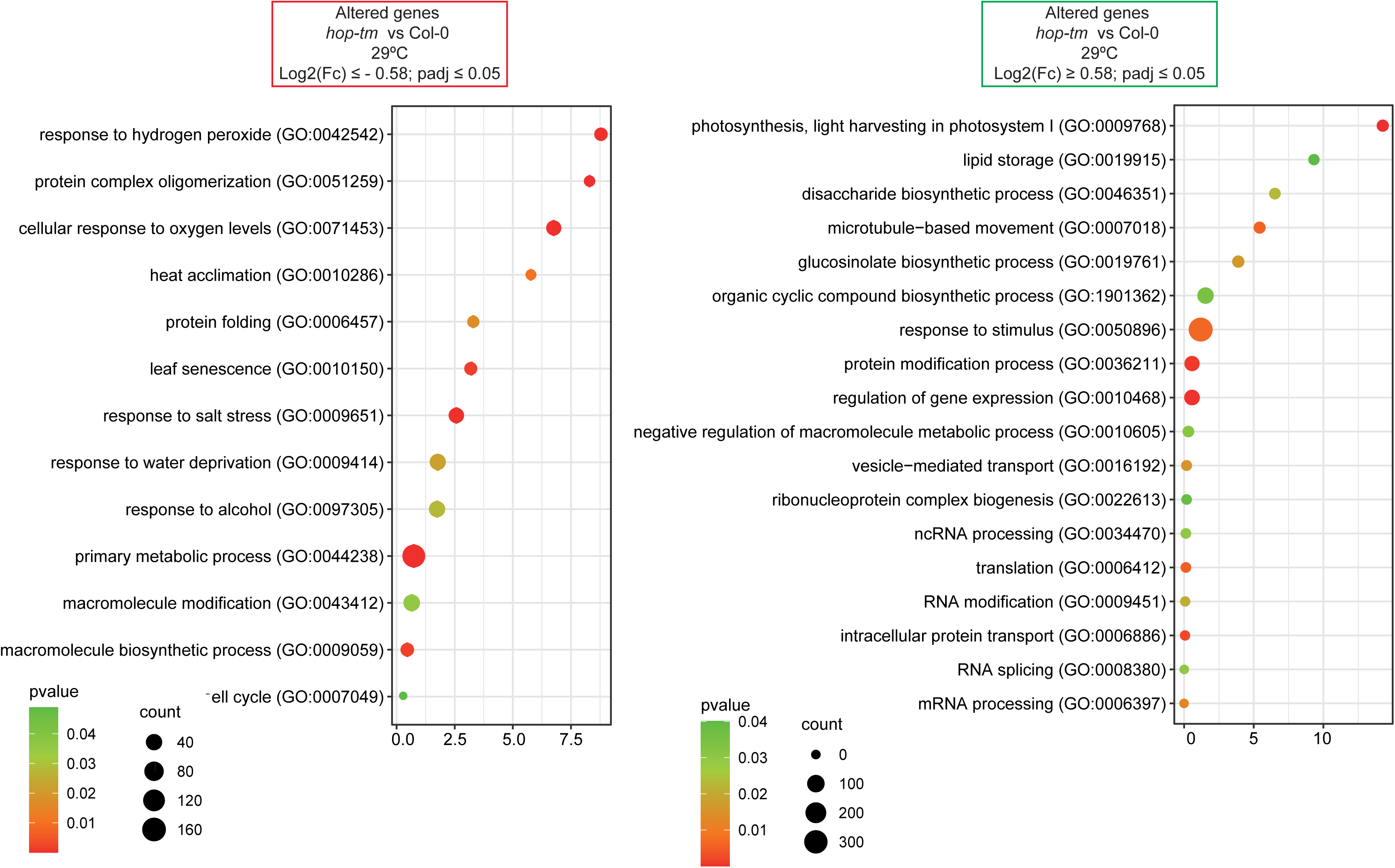
GO enrichment of the genes altered in the *hop1 hop2 hop3* mutant at warm temperature. Bubble plots showing the GO analysis of the significant downregulated genes (log2 (Fc) :: - 0.58; padj :: 0.05) (left panel) and upregulated genes (log2 (Fc) 2’ 0.58; padj :: 0.05) (right panel) in the *hop1 hop2 hop3* triple mutant (*hop-tm)* compared to Col-0 at 29°C.This analysis was done using Fisher statistical test and FDR for the analysis of p-value. Colour and count codes are shown at the right of the panel.

**Supplementary Figure 4.**
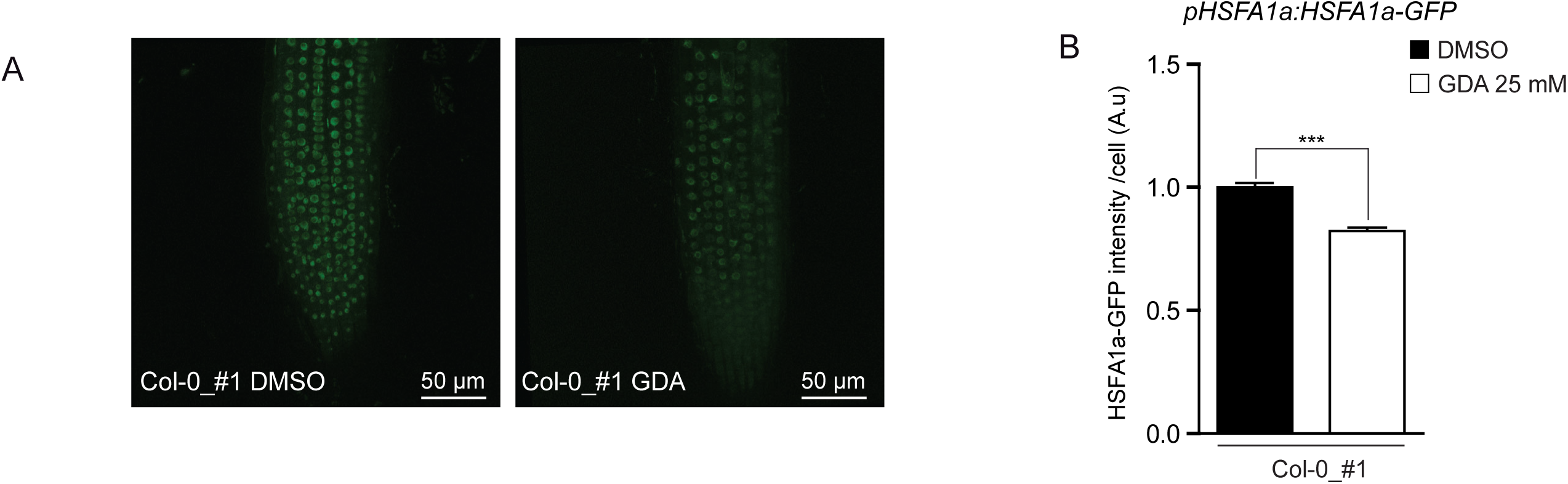
The stability of HSFA1a relies on the collaborative involvement of both HOP and HSP90. (A) Analysis of HSFA1a accumulation by confocal microscopy in 7-day-old seedling from the *pHSFA1a:HSFA1a-GFP* lines (Col-0 and hop-tm) of pair#1 treated in the absence or presence of 50 mM GDA for 5 h. Scale bars correspond to 50 µm. (B) quantification of GFP intensity per single cell nuclei n>1000 from the images and conditions described in (A). Statistically significant differences were calculated using Student’s t-test (ns, non-significant; *p < 0.05; **p < 0.01; ***p < 0.001). These experiments were performed three times obtaining similar results.

**Supplementary Table 1.**
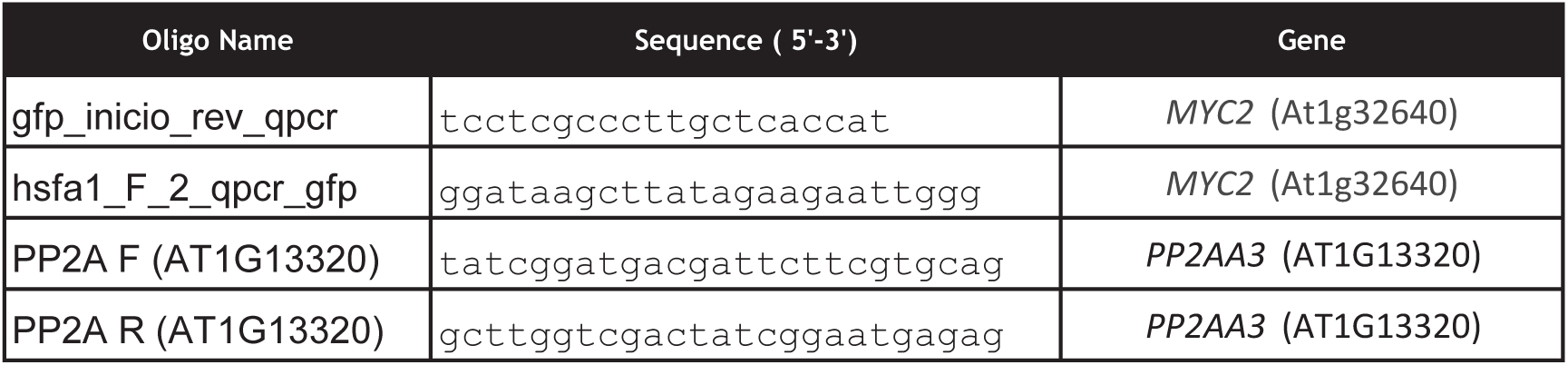
List of primers used in the study.

**Supplementary Table 2.** List of HSFA1a target genes that show an altered expression in the *hop1 hop2 hop3* (*hop_tm*) mutant compared to Col-0 at 29 °C.

| TAIR | SYMBOL | Log2 (Fc)<br>*hop-tm* vs Col-0<br>29°C | p<sub>adj</sub> |
| --- | --- | --- | --- |
| AT1G01560 | ATMPK11 | -5.4874 | 3.08E-20 |
| AT1G56060 | CYSTM3 | -4.4934 | 8.23E-05 |
| AT5G20230 | ATBCB | -3.5100 | 2.85E-26 |
| AT1G52890 | ANAC019 | -3.1625 | 3.98E-07 |
| AT2G41230 | ORS1 | -2.5127 | 5.58E-17 |
| AT1G76650 | CML38 | -2.3597 | 5.93E-10 |
| AT1G54050 | HSP17.4B | -2.2438 | 3.23E-33 |
| AT5G12030 | AT-HSP17.6A | -2.1284 | 6.50E-34 |
| AT1G07400 | HSP17.8 | -2.1142 | 7.22E-35 |
| AT2G38340 | DREB19 | -2.0396 | 1.33E-06 |
| AT1G66090 | TN3 | -2.0363 | 1.34E-06 |
| AT5G64750 | ABR1 | -1.8820 | 6.53E-08 |
| AT5G41740 | NA | -1.7792 | 1.91E-04 |
| AT2G18690 | NA | -1.7780 | 1.47E-03 |
| AT5G12020 | HSP17.6II | -1.6278 | 1.10E-17 |
| AT3G46230 | ATHSP17.4 | -1.6133 | 3.59E-43 |
| AT1G51090 | AtHMAD1 | -1.5574 | 3.20E-03 |
| AT4G12470 | AZI1 | -1.5253 | 1.19E-14 |
| AT4G02520 | ATGSTF2 | -1.5060 | 4.70E-07 |
| AT1G19020 | HUP35 | -1.4725 | 2.25E-13 |
| AT5G59820 | AtZAT12 | -1.3962 | 1.37E-14 |
| AT4G13395 | DVL10 | -1.3614 | 6.82E-04 |
| AT2G15080 | AtRLP19 | -1.3446 | 1.93E-02 |
| AT5G13210 | NA | -1.1928 | 1.44E-06 |
| AT1G17870 | ATEGY3 | -1.0774 | 4.53E-15 |
| AT1G18300 | atnudt4 | -1.0738 | 1.28E-06 |
| AT1G55530 | BTL06 | -0.9662 | 4.63E-16 |
| AT2G29500 | HSP17.6B | -0.9631 | 2.37E-15 |
| AT2G25140 | CLPB-M | -0.9529 | 4.41E-11 |
| AT2G26150 | HSFA2 | -0.9400 | 2.07E-05 |
| AT3G09350 | Fes1A | -0.9114 | 3.35E-10 |
| AT5G47830 | NA | -0.8967 | 6.68E-10 |
| AT2G46240 | ATBAG6 | -0.8804 | 3.69E-05 |
| AT1G66080 | HKL | -0.8795 | 1.10E-03 |
| AT5G52900 | MAKR6 | -0.8437 | 5.64E-07 |
| AT1G74310 | ATHSP101 | -0.8373 | 1.04E-09 |
| AT3G12400 | ATELC | -0.8231 | 6.36E-09 |
| AT3G09390 | ATMT-1 | -0.8183 | 2.76E-09 |
| AT1G19730 | ATH4 | -0.8042 | 1.87E-03 |
| AT3G48990 | AAE3 | -0.8029 | 5.21E-12 |

| **TAIR** | **SYMBOL** | **Log2 (Fc)**<br>***hop-tm* vs Col-0**<br>**29°C** | **padj** |
| --- | --- | --- | --- |
| AT4G37300 | MEE59 | -0.7985 | 4.07E-10 |
| AT1G59860 | HSP17.6A | -0.7910 | 6.05E-04 |
| AT5G14570 | ATNRT2.7 | -0.7897 | 9.86E-08 |
| AT1G21010 | NA | -0.7880 | 2.97E-06 |
| AT1G76070 | NA | -0.7855 | 3.26E-04 |
| AT5G35320 | NA | -0.7736 | 8.52E-07 |
| AT4G14430 | ATECI2 | -0.7680 | 1.92E-07 |
| AT2G23320 | AtWRKY15 | -0.7439 | 2.42E-09 |
| AT3G08970 | ATERDJ3A | -0.7436 | 5.08E-05 |
| AT5G02150 | Fes1C | -0.7288 | 5.45E-06 |
| AT3G20340 | NA | -0.7114 | 9.28E-05 |
| AT1G37130 | ATNR2 | -0.7083 | 9.50E-09 |
| AT5G46330 | FLS2 | -0.7021 | 4.59E-07 |
| AT5G51440 | HSP23.5 | -0.7018 | 1.12E-09 |
| AT1G71030 | ATMYBL2 | -0.6874 | 4.47E-05 |
| AT2G28400 | NA | -0.6761 | 2.42E-04 |
| AT3G60520 | NA | -0.6741 | 1.12E-02 |
| AT4G23570 | SGT1A | -0.6664 | 4.13E-07 |
| AT3G01560 | FLOE3 | -0.6621 | 5.02E-06 |
| AT5G62575 | SDH7 | -0.6545 | 2.48E-03 |
| AT2G36800 | DOGT1 | -0.6496 | 1.54E-05 |
| AT1G07410 | ATRAB-A2B | -0.6491 | 3.26E-02 |
| AT4G34540 | GVS1 | -0.6298 | 2.04E-03 |
| AT2G15960 | NA | -0.6096 | 6.18E-05 |
| AT5G14780 | FDH | -0.6042 | 5.55E-07 |
| AT4G11660 | AT-HSFB2B | -0.5979 | 2.53E-05 |
| AT4G24220 | 5[beta]-StR | -0.5942 | 4.97E-05 |
| AT5G43190 | NA | -0.5836 | 1.15E-03 |
| AT5G55180 | NA | 0.5815 | 7.32E-05 |
| AT3G59400 | GUN4 | 0.5823 | 4.87E-06 |
| AT1G78230 | NA | 0.5925 | 9.76E-05 |
| AT1G51400 | NA | 0.6037 | 2.75E-08 |
| AT4G05180 | PSBQ | 0.6174 | 3.28E-10 |
| AT4G34710 | ADC2 | 0.6214 | 8.92E-05 |
| AT5G64840 | ABCF5 | 0.6238 | 1.10E-03 |
| AT1G08380 | PSAO | 0.6241 | 4.69E-08 |
| AT4G02330 | AtPME41 | 0.6339 | 9.64E-03 |
| AT3G47470 | CAB4 | 0.6349 | 1.58E-08 |
| AT3G08940 | LHCB4.2 | 0.6481 | 3.74E-06 |
| AT4G23800 | 3xHMG-box2 | 0.6495 | 4.97E-02 |
| AT3G61060 | AtPP2-A13 | 0.6524 | 9.15E-08 |
| AT1G32060 | PRK | 0.6825 | 6.30E-12 |
| AT2G05100 | LHCB2 | 0.6827 | 5.68E-06 |
| AT2G31560 | NA | 0.6876 | 4.97E-05 |
| AT4G28750 | PSAE-1 | 0.7416 | 2.33E-10 |

| **TAIR** | **SYMBOL** | **Log2 (Fc)**<br>***hop-tm* vs Col-0**<br>**29°C** | **p<sub>adj</sub>** |
| --- | --- | --- | --- |
| AT3G26450 | NA | 0.7482 | 3.28E-09 |
| AT2G39730 | RCA | 0.7488 | 1.55E-15 |
| AT1G03130 | PSAD-2 | 0.7613 | 7.03E-10 |
| AT5G38430 | RBCS1B | 0.7718 | 3.79E-09 |
| AT1G30730 | AtBBE11 | 0.7912 | 2.84E-03 |
| AT1G52230 | PSAH-2 | 0.8060 | 1.46E-13 |
| AT4G27030 | FAD4 | 0.8289 | 1.04E-06 |
| AT5G19240 | NA | 0.8343 | 1.28E-05 |
| AT4G10340 | LHCB5 | 0.8408 | 3.10E-11 |
| AT4G33010 | AtGLDP1 | 1.1166 | 1.55E-24 |
| AT2G05070 | LHCB2 | 1.1402 | 3.69E-18 |
| AT3G02380 | ATCOL2 | 1.3715 | 3.73E-33 |
| AT5G54270 | LHCB3 | 1.4499 | 8.32E-29 |
| AT1G29910 | AB180 | 1.5720 | 6.02E-28 |
| AT1G29920 | AB165 | 1.7197 | 1.16E-30 |
| AT4G25140 | OLE1 | 3.6067 | 4.04E-03 |
| AT2G15020 | NA | 4.0287 | 3.21E-06 |
| AT2G42540 | COR15 | 4.0546 | 1.79E-05 |

